# The non-essentiality of essential genes suggests a loss-of-function therapeutic strategy for loss-of-function human diseases

**DOI:** 10.1101/040568

**Authors:** Piaopiao Chen, Dandan Wang, Han Chen, Zhenzhen Zhou, Xionglei He

**Affiliations:** State Key Laboratory of Biocontrol, School of Life Sciences, Sun Yat-sen University, Guangzhou 510275, China

**Author notes:** Correspondence to: Xionglei He, School of Life Sciences, Sun Yat-sen University, 135 Xingang West, Guangzhou 510275, China.

**Keywords:** Essential gene, Compensatory mutation, Conditional essentiality

## Abstract

Essential genes refer to those whose null mutation leads to lethality or sterility. We propose that the fatal effect of inactivating an essential gene can be attributed to either the loss of indispensable core cellular function (type I), or the gain of fatal side effects after losing dispensable periphery function (type II). In principle, inactivation of the type I essential genes can be rescued only by regain of the core functions, whereas inactivation of the type II essential genes could be rescued by a further loss of function of another gene to eliminate the otherwise fatal side effects. Because such loss-of-function rescuing mutations may occur spontaneously, type II essential genes may become non-essential in a few individuals of a large population. We tested this idea in the yeast *Sacchromyces cerevisiae*. Large-scale whole genome sequencing of such essentiality-reversing mutants reveals 14 cases where inactivation of an essential gene is rescued by loss-of-function mutations on another gene. In particular, the essential gene encoding the enzyme adenylosuccinate lyase (ADSL) is shown to be type II, suggesting a loss-of-function therapeutic strategy for the human disorder ADSL deficiency. A proof-of-principle test of this strategy in the nematode *Caenorhabditis elegans* shows promising results.

## Introduction

There are usually hundreds to thousands of essential genes in an organisms (Giaever et al. 2002; Kobayashi et al. 2003; Baba et al. 2006; Harris et al. 2010). It is often assumed that essential genes execute the functions that are indispensable to a cellular life(Mushegian and Koonin 1996; Koonin 2000). For example, in the bacterium *Escherichia coli* the essential gene DnaA is responsible for DNA replication initiation, TufA is responsible for RNA elongation in transcription, and InfA is responsible for translation initiation(Gerdes et al. 2003). The counterexamples, however, can be easily conceived. Suppose there are ten genes encoding a stable protein complex that carries out a dispensable function in a cell. Inactivation one of the ten genes may cause dosage imbalance, generating toxic intermediate that is lethal to the cell. Under this circumstance, each of the ten genes may appear essential while they together execute a dispensable function. Thus, there might be two types of essential genes: type I essential genes execute ‘core’ functions that are indispensable to the organism; type II essential genes execute ‘periphery’ functions that are dispensable to the organism, the lack of which, however, results in fatal side effects. Such a conceptual separation is meaningful, because inactivation of a type I essential gene can be rescued only by restoration of the core function, while inactivation of a type II essential gene could be rescued by further loss-of-function mutations on another gene to suppress the otherwise lethal side effects (Fig. 1). This reasoning has important implications to medical science because partial loss-of-function mutations on essential genes are involved in at least several hundred human inheritance disorders (Hamosh et al. 2005; Goh et al. 2007; Park et al. 2008). The prevailing therapeutic strategies for these diseases are to restore the lost functions (Garcia-Blanco et al. 2004; Maguire et al. 2008; Bidou et al. 2012). The proposition of type II essentiality suggests a possibility of using loss-of-function therapy to eliminate the disease-causing side effects in some disorders, which is conceptually much easier than the conventional gain-of-function therapy.

**Fig. 1.**
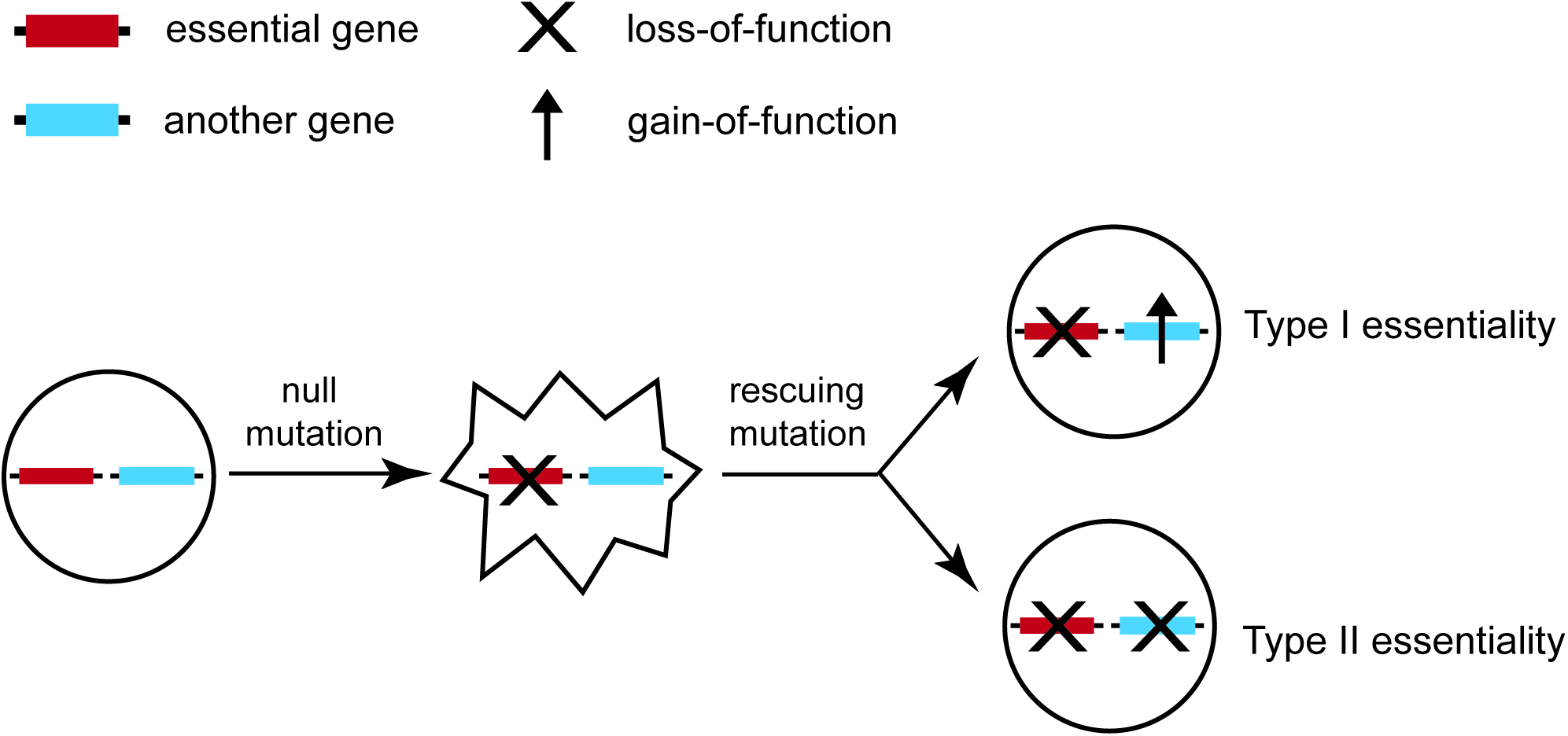
Two types of essential genes whose null phenotypes can be masked only by the gain of the original function (Type I) or by the loss of another function (Type II).

In this study we attempt to characterize type II essential genes using the model organism yeast *S. cerevisiae*. We reason that type II essential genes may become non-essential in a few individuals of a large population of the yeast cells since the potential rescuing mutations may occur spontaneously at a low frequency. By sequencing the genomes of the few individuals with such rescuing mutations we can identify the mutations that can mask the effects of inactivating the essential genes. Indeed, we obtained 17 such gene pairs where inactivation of an essential gene is rescued by loss-of-function mutations on the other gene, revealing a total of five type II essential yeast genes. Of particular interest is the gene encoding adenylosuccinate lyase, an enzyme in the purine *de novo* synthesis pathway. Partial loss-of-function mutations on this type II essential gene in humans cause adenylosuccinate lyase deficiency (ADSL; OMIM 103050), a rare Mendelian disorder with mental retardation and seizures as typical symptoms(Georges and Berghe 1984; Jaeken et al. 1988; Van den Berghe et al. 1997). We suggest a loss-of-function therapeutic strategy to suppress the phenotypes of ADSL deficiency. A proof-of-principle test of the strategy in the nematode *C. elegans* shows promising results.

## Results

### The rationale of using spontaneous mutations to reveal type II essentiality

We started with the yeast Tet-promoters Hughes Collection (yTHC) that is composed of 800 strains; in each strain the endogenous promoter of an essential gene is replaced by a tetracycline (TET) promoter(Mnaimneh et al. 2004). In principle, for a given strain the expression of the focal essential gene will be shut off upon the addition of doxycycline to the growth medium (Dox^+^ medium), resulting in no viable cell. However, if the focal essential gene is type II whose expression shut-off can be rescued by mutations on other genes, there might be some viable individuals among a large number of cells because such rescuing mutations can occur spontaneously despite at a low rate. In other words, we may simply plate a certain number of cells of an yTHC strain onto an agar plate supplied with doxycycline. Some viable clones will appear if the focal essential gene is type II.

We ran a computer simulation to estimate the number of cells required for capturing the spontaneous mutations that can mask the type II essentiality. We considered only critical loss-of-function mutations that include severe non-synonymous mutations, truncating substitution mutations and frame-shift indels. We started from a single cell and ended with a population of a given size *N*. The probability that at least one critical loss-of-function mutation occurs during the population expansion is estimated for each of the ~6,000 yeast genes (Methods). Up to ~84% of the genes can reach a probability of >90% when *N* = 3×10^7^, and no apparent increase is observed with larger *Ns* (Fig. 2A). The remaining genes with a lower probability of acquiring a critical loss-of-function mutation in the population of *N* = 3×10^7^ are mostly of short length (Fig. 2B), and it seems difficult to achieve a satisfactory detection probability for the ~15% yeast genes with a length of <800 base pairs. At any rate, there is a reasonably high probability to capture the potential spontaneous rescuing mutations if *N* = 3×10^7^ yeast cells are tested.

**Fig. 2.**
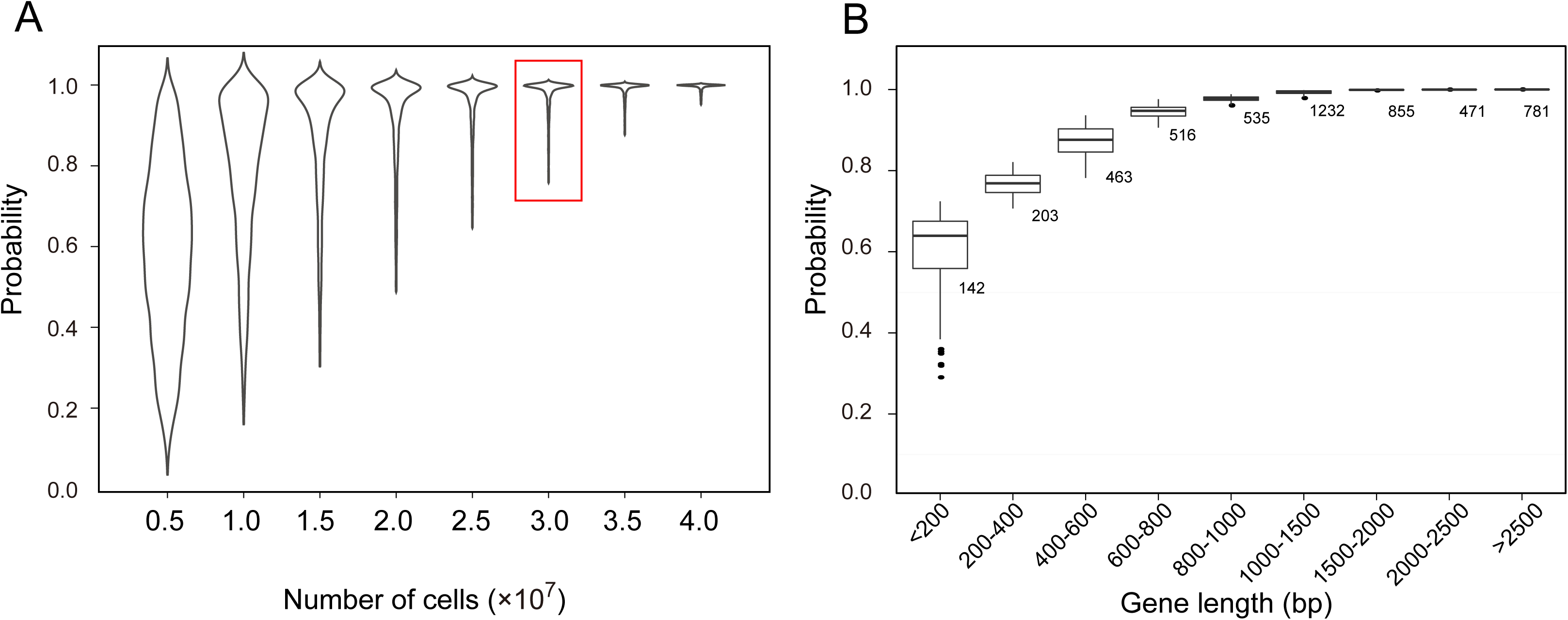
The probability of individual genes with at least one critical loss-of-function mutation during the population expansion from a single yeast cell. **(A)** The probability distribution of the ~6,000 yeast genes (y-axis) as a function of the ending population size *N* (x-axis). The probabilities have no apparent increase after the *N* reaches 3 × 10^7^. **(B)** The probability distribution (y-axis) as a function of the gene length (x-axis) when *N* = 3× 10^7^. The number of genes in each length category is shown next to the box.

### Characterization of type II essentiality

Among the 800 yTHC yeast strains some can grow well on the Dox^+^ medium. This is possibly because the TET promoter is not 100% shut-off, confounding the above strategy of revealing type II essential genes. To circumvent this problem, for each yTHC stain we first tested 10^4^~10^5^ cells and obtained 280 strains each showing no single colony on Dox^+^ agar. Only these strains are further examined. For each of the 280 strains we tested ~3×10^7^ cells on Dox^+^ agar plates and observed a few colonies in some plates (Methods) (Fig. 3A). There are two possibilities underlying the observation: mutations that invalidate the Tet-off system occurred such that the focal essential gene expresses normally on the Dox^+^ agar; alternatively, mutations that mask the focal gene essentiality occurred. In cases of the second possibility, the focal essential gene can be physically removed from the genomes of the clones grown on the Dox^+^ plates. We carried out the corresponding gene deletions using homologous recombination, and confirmed that, for five essential genes *ADE13, ERD2, MIM1, SEC14* and *RER2*, there are viable clones on the Dox^+^ plates explained by the second possibility.

**Fig. 3.**
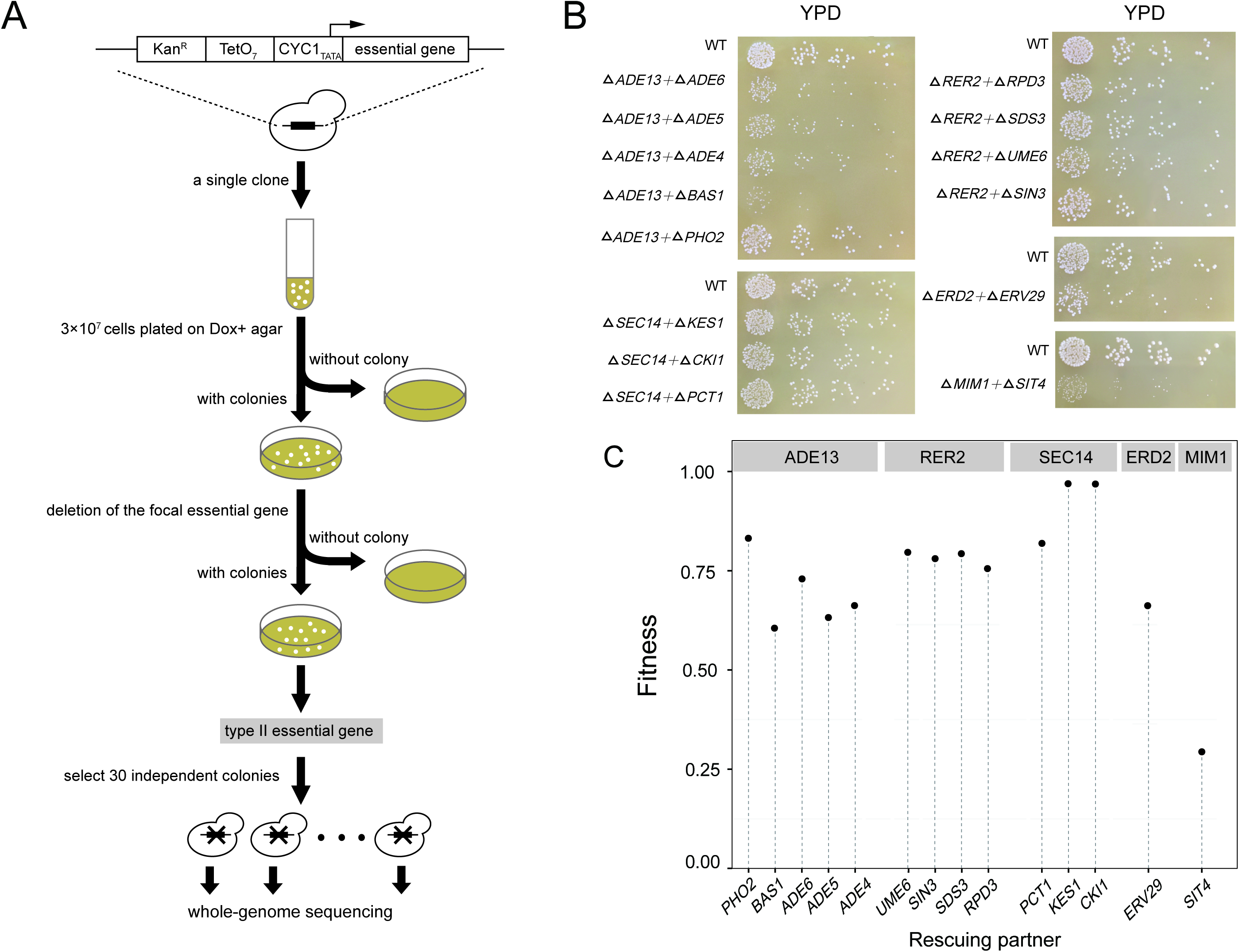
Identification of the type II essential genes and their rescuing partners. **(A)** A flowchart of the entire experimental procedure. **(B)** The colony size of each of the double deletion mutants. Cells diluted in gradient were spread on YPD plate, incubated for 48 hours except for the *ΔMIM1+ΔSIT4* that was incubated for 72 hours. Each panel shows a type II essential gene and its rescuing partners. **(C)** The fitness of each double gene deletion mutant relative to the wild-type in the rich medium YPD.

We focused on the five essential genes and expanded the plating experiment to obtain for each strain 30 largely independent clones in which the focal essential gene can be physically removed. To map the potential rescuing mutations we sequenced the genomes of the 30 x 5 = 150 clones and compared them with that of the parental strain of the yTHC stock. There are often quite a few *de novo* mutations found in a clone. For the 30 clones with the same focal essential gene, independent mutations found in the same gene(s) are likely to be the ones that can mask the focal gene essentiality. Indeed, we found a dozen of genes each enriched with the *de novo* mutations. Interestingly, these mutations appear to be mostly loss-of-function, suggesting that suppression of these genes may rescue the otherwise lethal effect upon silencing the focal essential gene, a phenomenon characteristic of type II essentiality (Table S1).

To test this we again carried out the corresponding gene deletions using homologous recombination. We started with BY4743, a diploid yeast strain with the wild-type genotype unless otherwise stated, and each time deleted one copy of the focal essential gene and one copy of the putative rescuing gene. Analysis of the haploid segregants of the double heterozygous deletion mutant help confirm whether the focal essential gene is indeed type II (Method). Overall, we obtained 14 such cases in which inactivating one of the five essential genes is rescued by inactivation of another gene that is invariably non-essential (Table 1). This result proves the five genes type II essential. The rescuing effects are generally strong since the growth rates of the double deletion mutants in the rich medium YPD are often over 70% of the wild-type rate (Fig. 3B and C).

**Table 1.**
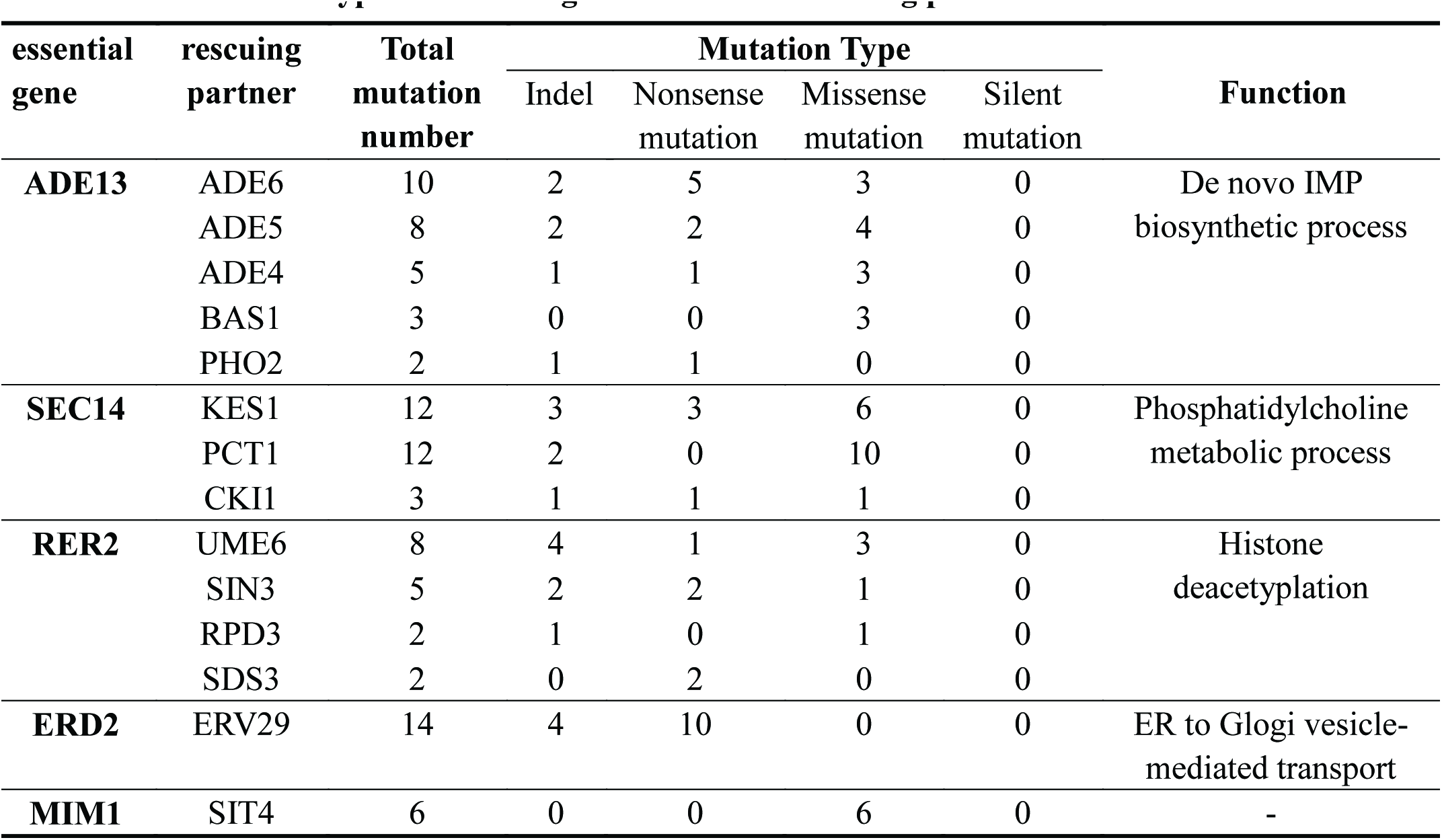
The identified type II essential genes with their rescuing partners.

| essential gene | rescuing partner | Total mutation number | Mutation Type |  |  |  | Function |
| --- | --- | --- | --- | --- | --- | --- | --- |
|  |  |  | Indel | Nonsense mutation | Missense mutation | Silent mutation |  |
| **ADE13** | ADE6 | 10 | 2 | 5 | 3 | 0 | De novo IMP biosynthetic process |
|  | ADE5 | 8 | 2 | 2 | 4 | 0 |  |
|  | ADE4 | 5 | 1 | 1 | 3 | 0 |  |
|  | BAS1 | 3 | 0 | 0 | 3 | 0 |  |
|  | PHO2 | 2 | 1 | 1 | 0 | 0 |  |
| **SEC14** | KES1 | 12 | 3 | 3 | 6 | 0 | Phosphatidylcholine metabolic process |
|  | PCT1 | 12 | 2 | 0 | 10 | 0 |  |
|  | CKI1 | 3 | 1 | 1 | 1 | 0 |  |
| **RER2** | UME6 | 8 | 4 | 1 | 3 | 0 | Histone deacetylation |
|  | SIN3 | 5 | 2 | 2 | 1 | 0 |  |
|  | RPD3 | 2 | 1 | 0 | 1 | 0 |  |
|  | SDS3 | 2 | 0 | 2 | 0 | 0 |  |
| **ERD2** | ERV29 | 14 | 4 | 10 | 0 | 0 | ER to Glogi vesicle-mediated transport |
| **MIM1** | SIT4 | 6 | 0 | 0 | 6 | 0 | - |

### Mechanisms underlying the type II essentiality

Gene Ontology (GO) analysis shows that the five type II essential genes and their rescuing partners often have the same GO terms (Table 1). For example, the essential gene *ERD2* and its non-essential partner *ERV29* are both involving in “Regulation of endoplasmic reticulum(ER) to Golgi vesicle-mediated transport”. *ERD2* is responsible for the recycling of vesicle-associated proteins between ER and Golgi and suppression of *ERD2* leads to abnormal Golgi structure due to the accumulation of too many proteins(Hardwick et al. 1992; Townsley et al. 1994). Interestingly, *ERV29* is responsible for transporting proteins from ER into Golgi. This suggests a likely mechanism of the *ERD2-ERV29* interaction: suppression of *ERD2* causes detrimental protein accumulation in Golgi, which is alleviated by suppressing *ERV29* to slow down the transportation of proteins into Golgi. In line with this, it has been shown that overexpression of *SED1-4*, four genes responsible for transporting proteins out of Golgi, can rescue the otherwise lethal effect of deleting *ERD2*(Hardwick et al. 1992).

A more intriguing case is the essential gene *ADE13*, which encodes the enzyme adenylosuccinate lyase (ADSL) and has five non-essential rescuing partners revealed by our screening. These genes are all related to *de novo* purine biosynthesis. A close examination of the *de novo* purine biosynthesis pathway shows that, with the exception of *ADE13*, all genes in the pathway are non-essential with negligible fitness reduction upon deletion(Giaever et al. 2002; Qian et al. 2012). This suggests that the process of *de novo* purine biosynthesis *per se* is dispensable to yeast cell in the rich medium YPD, so the essentiality of *ADE13* must be due to factors unrelated to purine production. We hypothesized that deletion of the enzyme gene results in accumulation of its substrates, which might be toxic to cell and thus fatal to the yeast. If this hypothesis is true, deletion of the upstream genes to block the production of S-AMP and SAICAR, the two substrates of *ADE13*, should be able to mask the effect of *ADE13* deletion (Fig. 4A).

**Fig. 4.**
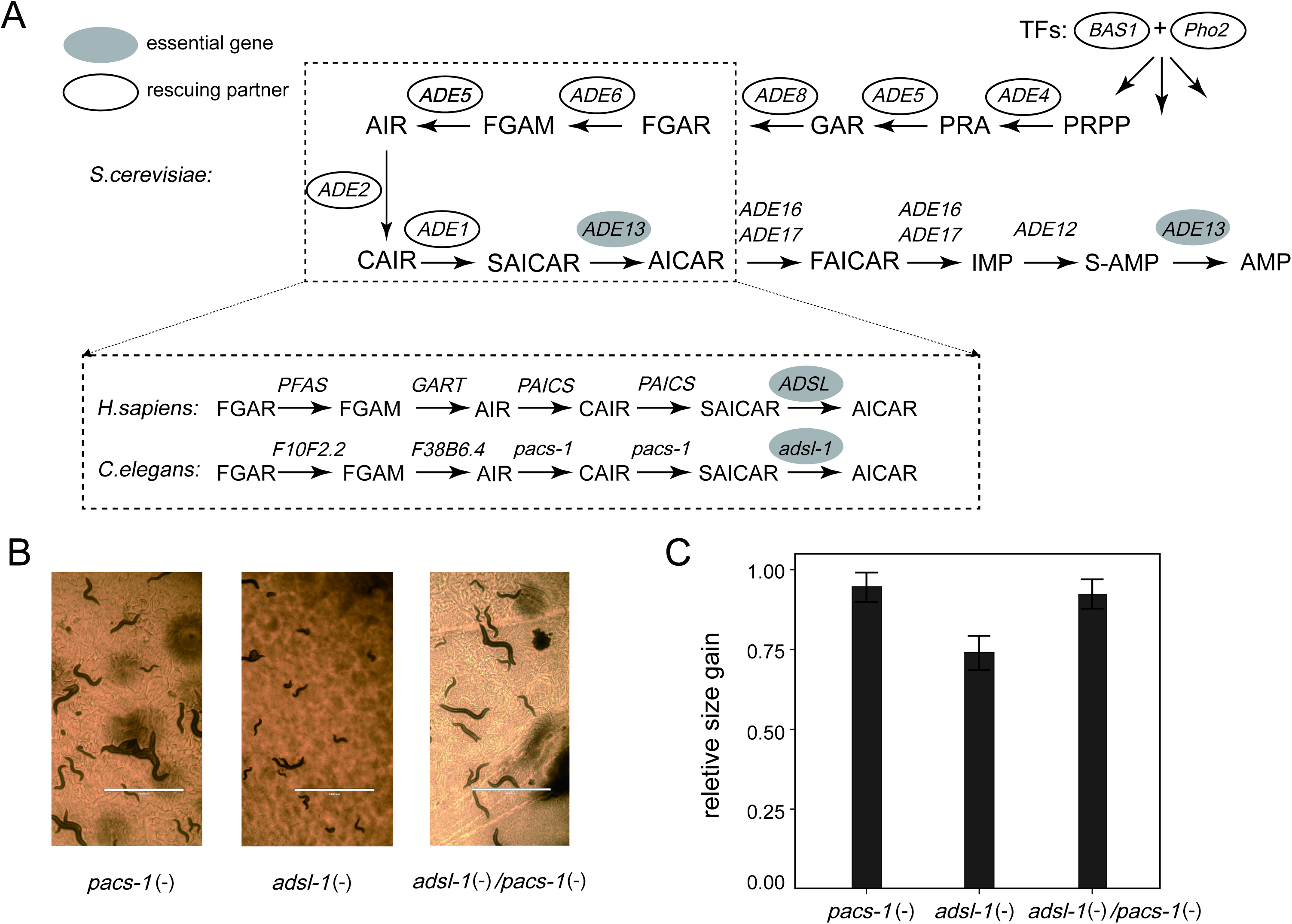
The loss-of-function therapy for the ADSL deficiency. **(A)** A schematic presentation of the purine *de novo* synthesis pathway. The pathway is extremely conserved among the yeast, nematode and humans. The filled oval marks the essential gene and the unfilled ovals show the rescuing partners of the essential gene. **(B)** The worms were fed with vectors producing the double-stranded RNAs each silencing *adsl-1, pacs-1* and *adsl-1*/*pacs-1*, respectively. Knockdown of *pacs-1* masks well the phenotypic defects of *adsl-1* knockdown. Scale bars shows 1000μm. **(C)** The relative size gain of the F1 worms from day-3 to day-6. The worms of *adsl-1 (-)/pacs-1(-)* have significantly faster growth than those of *adsl-1(-) (P* 10^−11^; t-test).

Consistent with this hypothesis, *ADE4, ADE5, ADE6, BAS1* and *PHO1*, the five rescuing partners revealed by our screening appear to be all upstream of *ADE13* in the pathway. Specifically, *ADE4, ADE5* and *ADE6* encode enzymes directly involved in the *de novo* purine biosynthesis and *BAS1* and *PHO1* are two transcription factors regulating the pathway(Denis et al. 1998). However, our screening provides no evidence that the other three upstream genes in the pathway, namely *ADE8, ADE2* and *ADE1*, can mask the effect of *ADE13* deletion. We thus performed homologous recombination-based gene deletions and found that deleting each of the three genes is able to mask the otherwise lethal effect of *ADE13* deletion. This result well supports the above substrate accumulation hypothesis for explaining the type II essentiality of *ADE13. ADE13* catalyzes two reactions with S-AMP and SAICAR as substrates, respectively (Van den Bergh et al. 1993). It is unclear which one is responsible for the lethality of *ADE13* deletion. If S-AMP is the toxin, deletion of *ADE12*, the enzyme gene directly responsible for its production, should mask the effect of *ADE13* deletion. Alternatively, deletion of *ADE1*, the gene encoding the enzyme phosphoribosylaminoimidazole carboxylase (PAICS), should mask the effect of *ADE13* deletion if SAICAR is the toxin. We successfully obtained *Δade13+Δade1* double mutant but failed to obtain *Δade13+Δade12* double mutant, suggesting that the accumulation of SAICAR, rather than S-AMP, is the cause of the lethal effect upon *ADE13* deletion.

### Loss-of-function therapy for the loss-of-function ADSL disease

The purine *de novo* biosynthesis pathway is conserved among various lineages including yeast, nematode and mammals(Kanehisa and Goto 2000) (Fig. 4A). Importantly, partial loss-of-function mutations on the human gene encoding the enzyme ADSL cause ADSL deficiency, an ultra-rare Mendelian disease with mental retardation and seizures as typical symptoms(Georges and Berghe 1984; Jaeken et al. 1988; Van den Berghe et al. 1997). To date there is no treatment with proven clinical efficacy for this disease despite a variety of therapeutic attempts (Jaeken et al. 1988; Salerno et al. 1998; Ciardo et al. 2001; Salerno et al. 2002; Jurecka et al. 2008). For such loss-of-function diseases, gain-of-function therapeutic strategy is predominant. Interestingly, recognition of the type II essentiality of the gene encoding ADSL suggests a loss-of-function therapeutic strategy to suppress the phenotypes of ADSL deficiency, which is conceptually much easier than the conventional gain-of-function strategy to restore the lost function. We decided to test this idea in the nematode *C. elegans.* The nematode genes encoding the two enzymes *ADSL* and *PAICS* are *adsl-1* and *pacs-1*, respectively. RNA interference (RNAi) is used to mimic the loss of function of the two genes in *C. elegans* (Methods).

Body length is used to assess the effect upon knocking down the *C. elegans* genes. We considered the average increase of body length of a worm from the day 3 to day 6 after its birth. As expected, knocking down the essential *adsl-1* encoding gene alone results in a substantial reduction of the body growth compared to the negative control, while knocking down the non-essential *pacs-1* encoding gene shows no significant difference from the negative control (Fig. 4B and C). Interestingly, the effect of knocking down simultaneously the two genes is largely the same as that of knocking down the *pacs-1* encoding gene alone. This result indicates that the phenotype of ADSL deficiency in the nematode *C. elegans* is nearly fully masked by a further loss-of-function perturbation on the gene encoding *pacs-1*. Given the identical purine de novo biosynthesis pathway between the *C. elegans* and humans, the loss-of-function therapeutic strategy proposed for the ADSL deficiency is highly likely to be effective in humans.

## Discussion

In this study, we propose two types of gene essentiality: the type I can be rescued/masked only by gain-of-function perturbations while the type II can be rescued/masked by loss-of-function perturbations. This proposition is different from the previously proposed concept of genetically conditional essentiality that states nothing about the way of rescuing (Zhao et al. 1998; Geissler et al. 2003; Gerdes et al. 2006; Rancati et al. 2008; Vernon et al. 2008; Delia et al. 2009; Bergmiller et al. 2012). Taking advantages of spontaneous mutations, a useful tool for probing the gene network(Szamecz et al. 2014; Liu et al. 2015), as well as the affordable whole genome sequencing, we identified five type II essential genes each with 1-8 rescuing partners in the yeast. This is a preliminary screening and there are good reasons to expect many more type II essential genes: First, only one fourth of the yeast essential genes are examined because of the leakiness of the Tet promoter, and for each gene only ~3×10^7^ cells are tested. We found that plating more cells produces viable colonies that are primarily due to the failure of the TET system as a result of spontaneous mutations. Second, the mutation rate varies among genes by orders of magnitude, so our strategy that is designed based on the average mutation rate becomes invalid for detecting the rescuing mutations on the genes with intrinsically low mutation rate. This point is well demonstrated in the case of *ADE13* that has eight rescuing partners; our extensive screening identifies only five of the eight and the rest three are suggested by reasoning and validated by target gene deletions. Third, a recent study reports that up to ~10% of the yeast essential genes are dispensable in a newly evolved genetic status(Liu et al. 2015). Although the resulting aneuploidy precludes the authors from identifying the specific genes responsible for the essentiality turnover, it won’t be surprising if a further study shows a large proportion of them type II essential. Interestingly, only one of the five type II essential genes identified in this study is found conditionally dispensable in that study, suggesting that both screenings are far from saturation.

It is worth noting that such type II essentiality was actually observed before in quite a few cases, including the yeast gene *ADE13*(Zekhnov et al. 1995; Zekhnov and Domkin 2000), although the knowledge has never been applied to thinking of the therapy for the human disease. The clinical difficulty for Mendelian diseases is in great part due to the fact that most of the diseases are due to loss-of-function mutations on the genes with key functions. The basic therapeutic principle for such diseases has been to restore the lost functions (Garcia-Blanco et al. 2004; Maguire et al. 2008; Bidou et al. 2012), which is extremely difficult to realize. Because inventing a compound to inactive a function is much simpler than inventing a compound to restore a function, using loss-of-function perturbations to mask the loss-of-function effects of type II essential genes suggests a conceptually much easier therapeutic strategy for loss-of-function human diseases. Therefore, studies of the nature of essential genes not only deepen our understandings of the gene network but can also help the medical practice.

## Methods

### Yeast strains

The Tet promoter-based Hughes Collection (yTHC) of yeast strains were purchased from GE Healthcare Dharmacon. The kanR-TetO7-TATA cassette was integrated into the yeast genome, replacing the endogenous promoter for each essential gene in the haploid background strain R1158 *(URA3∷CMV-tTA MATa his3-1 leu2-0 met15-0)*. This strain was created by a one-step integration of the tet-off activator (tTA, which dissociated in the presence of doxycycline), under the control of the CMV promoter, at the URA3 locus. The diploid yeast strain used in the study is BY4743 (*MATa/α his3Δ1/ his3Δ1 leu2Δ0/ leu2Δ0 lys2Δ0/ lys2Δ0 metΔ15/met15 ura3Δ0 /ura3Δ0*).

### Simulating the number of cells required for capture the rescuing mutations

In the previous study, an overall base-substitutional mutation rate and small insertion/deletions mutation rate estimate of 0.33 × 10^−9^ and 0.02× 10^−9^ per site per cell division(Lynch et al. 2008). Analysis of the loss-of-function mutation probability for each gene must consider the severe non-synonymous mutations, truncating substitution mutations and frame-shift indels. The probability is estimated that at least one null mutation occurs during the population expansion for each of the ~6,000 yeast genes when screened different population size *Ns.* The probability that occur loss-of-function mutation for each gene is estimated by

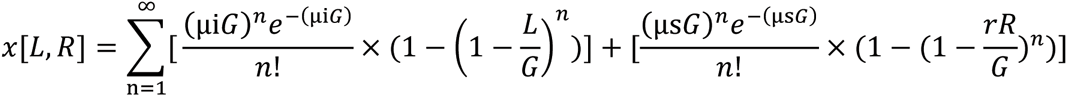

Where μ_i_ is the indel mutation rate per base per generation; μ_s_ is the single-nucleotide mutation rate per base per generation(Lynch et al. 2008). L is the gene length while G represent the whole genome size. R represent the number of non-synonyms substitutions sites while r is the ratio of severe sites upon all non-synonyms substitutions sites. For a given population *N*, the probability was estimated by 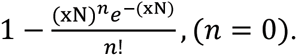

### Screening assay

Yeast Tet-promoter strains for screening were removed from frozen stock (25% glycerol, −80 °C) and streaked onto YPD plate (1%Yeast Extract, 2%Peptone, 2%Dextrose). A single original colony were inoculated in YPD medium overnight. For each strain, 3×10^7^ cells were condensed, washed and then spread on YPD plate with 10 *μ*g/ml doxycycline, incubated for 3-4 days at 30°C. On some Dox^+^ plates, there were indeed appeared some viable colonies, and then we attempted to knock down the focal essential genes of these viable colonies using standard lithium acetate transformation protocol. We successfully detected positive deletion in five essential genes. For each of the five genes, the screening experiment mentioned above were repeated 30 times to select 30 independent clones on 30 Dox^+^ plates. Finally, 150 (30×5) clones were all deleted focal essential gene to exclude the false positive mutation. By the way, the five parental clones were prepared to sequence as well.

### Genome sequencing and mutation calling

Genomic DNA was extracted from ~10^8^ yeast cells using the Dr. GenTLE® (from Yeast) High Recovery kit (catalog number: 9082). Three genomes were pooled together and we had 55(10×5+5) samples for the five essential genes. Fragments between 400bp and 600bp were collected for library construction and samples were sequenced on an Illumina Hi-seq 2000 platform. Approximately 8 million 125bp reads were generated for each sample, corresponding to an average sequencing depth of ~80. Sequence reads were aligned to yeast genome by bowtie2(Langmead and Salzberg 2012) with default setting, and duplicated reads were removed by Picard (http://picard.sourceforge.net). Single-nucleotide mutations and indels were called on the Genome Analysis Toolkit (GATK) platform(Margulies et al. 2005) with default settings.

### Identification of the type II essential genes

The gene Leu and Ura were used to replace one copy of essential gene and their rescuing gene respectively in diploid yeast strain BY4743 using a standard lithium acetate transformation protocol. The resulting double heterozygous deletion mutant strains were inoculated in sporulation medium (10g/L potassium acetate and 50mg/L zinc acetate in water) for 4-5 days at 25°C on a rotary shaker. 200*μ*l of cell suspension were resuspended in Potassium Phosphate buffer (67 mM KH2PO4; pH 7.5) added with 25 units Zymolyase (ZYMO research), incubated for 30min at 30°C on a rotary shaker to digest the cell walls and then held in a dry bath for 15min at 55°C to kill diploid cells. Then, the spore suspension was spread on YPD plate for three days. Spores from the diploid strains heterozygous for a focal essential gene and a putative rescuing gene deletion would be expected to show a 1:1:1:0 viability segregation pattern between the four haploid spores: wild-type; rescuing gene deletion; both essential gene and rescuing gene deletion; only essential gene deletion. Analysis of the haploid segregants help confirm whether the focal essential gene is indeed type II.

### *C.elegans* culture and gene knockdown using RNAi

*C.elegans* strains (rrf-3-/-) were maintained at 16°C on NGM plate, with *E.coli* (OP50) as feed (WormBook). We use RNA interference (RNAi) to mimic loss-of-function mutation of gene *adsl-1* and *pacs-1* in C. elegans. Two single-gene-interferencing vectors, *adsl-1*/GFP and *pacs-1*/GFP, for knocking down *adsl-1* and *pacs-1* separately and a double-gene-interferencing vector, *adsl-1*/*pacs-1*, for knocking down the two genes simultaneously, were constructed. We fused the gene with the coding sequence of GFP in the single-gene-interferencing vectors to achieve a similar length of insert as the double-gene-interferencing vector, which ensures comparable RNAi efficiency for the same genes in different vectors. The L4440 empty vector were used as negative control. The L4440 vector and the L4440 with gene of interest vectors were transformed into the HT115 strain, then grown in LB media with ampicillin at 37°C. The next day, bacteria were 1:100 transferred into fresh LB media at 16°C and used 1mM IPTG to induce dsRNA expression. The bacteria were harvested and transferred onto NGM plates containing IPTG as feed. For each RNAi experiment, at least ten three-day-old F0 worms were transferred onto an RNAi plate. The F0 worms were killed with heated inoculating loop at day 0, when F1 eggs were first observed on the plate. Growth of F1 worms were measured in day 3 and day 6, using ImageJ.

### Quantitative PCR

Total RNA was extracted from the *C. elegans* worms using Trizol (Life Technologies, catalog number: 15596-026). Two micrograms of total RNA were treated by RNase-Free DNase Set (QIAGEN, catalog number: 79254) and converted into cDNA using PrimerScript RT reagent Kit (Takara, catalog number: RR037A). The qPCR experiments were performed using SYBR Green (Takara, catalog number: RR820B) and Roche LightCycler 480 apparatus. The *β*-actin gene was used as the reference gene. Quantitative RT-PCR reveals successful RNA interference. Expression level of the gene *adsl-1* reduces by 64.4% and 61.1% in worms subject to *adsl-1*/GFP and *adsl-1*/*pacs-1* treatment, respectively; the numbers are 62.1% and 58.2% for the gene *pacs-1* in worms subject to *pacs-1*/GFP and *adsl-1*/*pacs-1* treatment, respectively.

## Acknowledgements

The *C. elegans* and *E. coli* strains used in the RNAi experiment are gifts of Dr. X. Wang of NIBS.

## References

Baba T, Ara T, Hasegawa M, Takai Y, Okumura Y, Baba M, Datsenko KA, Tomita M, Wanner BL, Mori H. 2006. Construction of Escherichia coli K-12 in-frame, single-gene knockout mutants: the Keio collection. Molecular systems biology 2(1).

Bergmiller T, Ackermann M, Silander OK. 2012. Patterns of evolutionary conservation of essential genes correlate with their compensability.

Bidou L, Allamand V, Rousset J-P, Namy O. 2012. Sense from nonsense: therapies for premature stop codon diseases. Trends in molecular medicine 18(11): 679–688.

Ciardo F, Salerno C, Curatolo P. 2001. Topical Review: Neurologic Aspects of Adenylosuccinate Lyase Deficiency. Journal of Child Neurology 16(5): 301–308.

Delia MA, Pereira MP, Brown ED. 2009. Are essential genes really essential? Trends in Microbiology 17(10): 433–438.

Denis V, Boucherie H, Monribot C, Daignan-Fornier B. 1998. Role of the myb-like protein bas1p in Saccharomyces cerevisiae: a proteome analysis. Molecular Microbiology 30(3): 557–566.

Garcia-Blanco MA, Baraniak AP, Lasda EL. 2004. Alternative splicing in disease and therapy. Nature biotechnology 22(5):535–546.

Geissler B, Elraheb D, Margolin W. 2003. A gain-of-function mutation in ftsA bypasses the requirement for the essential cell division gene zipA in Escherichia coli. Proceedings of the National Academy of Sciences 100(7): 4197–4202.

Georges J, Berghe V. 1984. An infantile autistic syndrome characterised by the presence of succinylpurines in body fluids. The Lancet 324(8411): 1058–1061.

Gerdes S, Edwards R, Kubal M, Fonstein M, Stevens R, Osterman A. 2006. Essential genes on metabolic maps. Current Opinion in Biotechnology 17(5): 448–456.

Gerdes S, Scholle M, Campbell J, Balazsi G, Ravasz E, Daugherty M, Somera A, Kyrpides N, Anderson I, Gelfand M. 2003. Experimental determination and system level analysis of essential genes in Escherichia coli MG1655. Journal of bacteriology 185(19): 5673–5684.

Giaever G, Chu AM, Ni L, Connelly C, Riles L, Veronneau S, Dow S, Lucau-Danila A, Anderson K, Andre B. 2002. Functional profiling of the Saccharomyces cerevisiae genome. nature 418(6896): 387–391.

Goh K-I, Cusick ME, Valle D, Childs B, Vidal M, Barabasi A-L. 2007. The human disease network. Proceedings of the National Academy of Sciences 104(21): 8685–8690.

Hamosh A, Scott AF, Amberger JS, Bocchini CA, McKusick VA. 2005. Online Mendelian Inheritance in Man (OMIM), a knowledgebase of human genes and genetic disorders. Nucleic acids research 33(suppl 1): D514–D517.

Hardwick KG, Boothroyd JC, Rudner AD, Pelham H. 1992. Genes that allow yeast cells to grow in the absence of the HDEL receptor. The EMBO journal 11(11): 4187.

Harris TW, Antoshechkin I, Bieri T, Blasiar D, Chan J, Chen WJ, De La Cruz N, Davis P, Duesbury M, Fang R. 2010. WormBase: a comprehensive resource for nematode research. Nucleic acids research 38(suppl 1): D463–D467.

Jaeken J, Wadman S, Duran M, Van Sprang F, Beemer F, Holl R, Theunissen P, De Cock P, Van den Bergh F, Vincent M-F. 1988. Adenylosuccinase deficiency: an inborn error of purine nucleotide synthesis. European journal of pediatrics 148(2): 126–131.

Jurecka A, Tylki-Szymanska A, Zikanova M, Krijt J, Kmoch S. 2008. D-ribose therapy in four Polish patients with adenylosuccinate lyase deficiency: Absence of positive effect. Journal of inherited metabolic disease 31(2): 329–332.

Kanehisa M, Goto S. 2000. KEGG: kyoto encyclopedia of genes and genomes. Nucleic acids research 28(1): 27–30.

Kobayashi K, Ehrlich SD, Albertini A, Amati G, Andersen K, Arnaud M, Asai K, Ashikaga S, Aymerich S, Bessieres P. 2003. Essential Bacillus subtilis genes. Proceedings of the National Academy of Sciences 100(8): 4678–4683.

Koonin EV. 2000. How Many Genes Can Make a Cell: The Minimal-Gene-Set Concept 1. Annual review of genomics and human genetics 1(1): 99–116.

Langmead B, Salzberg SL. 2012. Fast gapped-read alignment with Bowtie 2. Nature methods 9(4): 357–359.

Liu G, Yong MYJ, Yurieva M, Srinivasan KG, Liu J, Lim JSY, Poidinger M, Wright GD, Zolezzi F, Choi H. 2015. Gene Essentiality Is a Quantitative Property Linked to Cellular Evolvability. Cell.

Lynch M, Sung W, Morris K, Coffey N, Landry CR, Dopman EB, Dickinson WJ, Okamoto K, Kulkarni S, Hartl DL. 2008. A genome-wide view of the spectrum of spontaneous mutations in yeast. Proceedings of the National Academy of Sciences 105(27): 9272–9277.

Maguire AM, Simonelli F, Pierce EA, Pugh Jr EN, Mingozzi F, Bennicelli J, Banfi S, Marshall KA, Testa F, Surace EM. 2008. Safety and efficacy of gene transfer for Leber's congenital amaurosis. New England Journal of Medicine 358(21):2240–2248.

Margulies M, Egholm M, Altman WE, Attiya S, Bader JS, Bemben LA, Berka J, Braverman MS, Chen Y-J, Chen Z. 2005. Genome sequencing in microfabricated high-density picolitre reactors. Nature 437(7057): 376–380.

Mnaimneh S, Davierwala AP, Haynes J, Moffat J, Peng W-T, Zhang W, Yang X, Pootoolal J, Chua G, Lopez A. 2004. Exploration of essential gene functions via titratable promoter alleles. Cell 118(1): 31–44.

Mushegian AR, Koonin EV. 1996. A minimal gene set for cellular life derived by comparison of complete bacterial genomes. Proceedings of the National Academy of Sciences 93(19): 10268–10273.

Park D, Park J, Park SG, Park T, Choi SS. 2008. Analysis of human disease genes in the context of gene essentiality. Genomics 92(6): 414–418.

Qian W, Ma D, Xiao C, Wang Z, Zhang J. 2012. The genomic landscape and evolutionary resolution of antagonistic pleiotropy in yeast. Cell reports 2(5): 1399–1410.

Rancati G, Pavelka N, Fleharty B, Noll A, Trimble R, Walton K, Perera A, Staehling-Hampton K, Seidel CW, Li R. 2008. Aneuploidy underlies rapid adaptive evolution of yeast cells deprived of a conserved cytokinesis motor. Cell 135(5): 879–893.

Salerno C, Celli M, Finocchiaro R, D’Eufemia P, Iannetti P, Crifo C, Giardini O. 1998. Effect of D-ribose administration to a patient with inherited deficit of adenylosuccinase. In Purine and Pyrimidine Metabolism in Man IX, pp. 177–180. Springer.

Salerno C, Crifo C, Curatolo P, Ciardo F. 2002. Effect of uridine administration to a patient with adenylosuccinate lyase deficiency. In Purine and Pyrimidine Metabolism in Man X, pp. 75–78. Springer.

Szamecz B, Boross G, Kalapis D, Kovács K, Fekete G, Farkas Z, Lázár V, Hrtyan M, Kemmeren P, Koerkamp MJG. 2014. The genomic landscape of compensatory evolution.

Townsley FM, Frigerio G, Pelham H. 1994. Retrieval of HDEL proteins is required for growth of yeast cells. The Journal of cell biology 127(1): 21–28.

Van den Bergh F, Vincent M-F, Jaeken J, Van den Berghe G. 1993. Residual adenylosuccinase activities in fibroblasts of adenylosuccinase-deficient children: parallel deficiency with adenylosuccinate and succinyl-AICAR in profoundly retarded patients and non-parallel deficiency in a mildly retarded girl. Journal of inherited metabolic disease 16(2): 415–424.

Van den Berghe G, Vincent M-F, Jaeken J. 1997. Inborn errors of the purine nucleotide cycle: adenylosuccinase deficiency. Journal of inherited metabolic disease 20(2): 193–202.

Vernon M, Lobachev K, Petes TD. 2008. High rates of "unselected" aneuploidy and chromosome rearrangements in tel1 mec1 haploid yeast strains. Genetics 179(1): 237–247.

Zekhnov AM, Domkin VD. 2000. [The phenomenon of predetermination of the cytoplasm upon interaction between alleles of the ADE2 and ADE13 in the yeast Saccharomyces cerevisiae]. Genetika 36(4).

Zekhnov AM, Domkin VD, Démbéréliĭn O, Shubochkina EA, Smirnov MN. 1995. [Mutation of ade13-1 of the yeast Saccharomyces cerevisiae leads to the absence of growth on a complete medium with glucose and epistatically interacts with mutations in other genes for purine biosynthesis]. Genetika 31(1).

Zhao X, Muller EG, Rothstein R. 1998. A suppressor of two essential checkpoint genes identifies a novel protein that negatively affects dNTP pools. Molecular cell 2(3): 329–340.

